# Identification and functional characterisation of a locus for target site integration in *Fusarium graminearum*

**DOI:** 10.1101/2023.08.20.553861

**Authors:** Martin Darino, Martin Urban, Navneet Kaur, Ana Machado-Wood, Michael Grimwade-Mann, Dan Smith, Andrew Beacham, Kim Hammond-Kosack

## Abstract

**Background:** Fusarium Head Blight is a destructive floral disease of different cereal crops. The Ascomycete fungus *Fusarium graminearum* (*Fg*) is one of the main causal agents of FHB in wheat and barley. The role(s) in virulence of *Fg* genes include genetic studies that involve the transformation of the fungus with different expression cassettes. We have observed in several studies where *Fg* genes functions were characterised that integration of expression cassettes occurred randomly. Random insertion of a cassette may disrupt gene expression and/or protein functions and hence the overall conclusion of the study. Target site integration (TSI) is an approach that consists in identifying a chromosomal region where the cassette can be inserted. The identification of a suitable locus for TSI in *Fg* would avert the potential risks of ectopic integration.

**Results:** Here, we identified a highly conserved intergenic region on chromosome 1 suitable for TSI. We named this intergenic region the TSI locus 1. We developed an efficient cloning vector system based on the Golden Gate method to clone different expression cassettes for use in combination with TSI locus 1. We present evidence that integrations in the TSI locus 1 affects neither fungal virulence nor fungal growth under different stress conditions. Integrations at the TSI locus 1 resulted in the expression of different gene fusions. In addition, the activities of *Fg* native promoters were not altered by integration into the TSI locus 1. We have developed a bespoke bioinformatic pipeline to analyse the existence of ectopic integrations and tandem insertions of the cassette that may occurred during the transformation process. Finally, we established a protocol to study protein secretion in wheat coleoptiles using confocal microscopy and the TSI locus 1.

**Conclusion:** The TSI locus 1 can be used in *Fg* and potentially other cereal infecting Fusarium species for diverse studies including promoter activity analysis, secretion, protein localisation studies and gene complementation. The bespoke bioinformatic pipeline developed in this work can be an alternative to southern blotting, the gold standard technique to identify ectopic integration and tandem insertions in fungal transformation.

## Background

Fusarium Head Blight (FHB) is a destructive floral disease of different cereal crops such as wheat, barley, maize and oat (1, 2). The Ascomycete fungus *Fusarium graminearum* (*Fg*) is one of the main causal agents of FHB in wheat and barley crops in Europe, Asia and America (3). The disease is characterised by reducing grain quality and safety. During infection, *Fg* produces a diverse repertoire of mycotoxins where deoxynivalenol is one of the most frequent detected in cereal grains (4). Contamination of grains with different mycotoxins make the crop unsuitable for human and/or animal consumption (4). Due to the ever growing worldwide economic and societal relevance of FHB disease, the role(s) in virulence of 1571 *Fg* genes, i.e. 11 % of the predicted *Fg* gene repertoire, has been formally tested, described in different peer reviewed studies and then manually curated into the multispecies PHI-base database (5). These studies often include approaches such as gene deletion, gene complementation, promoter expression and protein localisation to characterise a gene and/or a protein function (6–8). Gene complementation and protein localisation involve the stable transformation of *Fg* with an expression cassette. Integration of the cassette into the genome can occur by either homologous or non-homologous recombination (6, 9). Non-homologous recombination or ectopic integration happens when the cassette is not flanked by sequences with homology to the native chromosomal region. Random insertion of a cassette may disrupt gene expression and/or protein functions and hence the overall conclusion of the study. Target site integration (TSI) is not a new concept in fungal molecular genetic studies. TSI has been described for different plant pathogens such as *Ustilago maydis* (*U. maydis*) and *Magnaporthe oryzae* (*M. oryzae*) (10, 11). The approach consists in identifying a chromosomal region where the cassette can be inserted by homologous recombination. A suitable locus for TSI is defined as a region where insertion of an expression cassette does not alter the growth and virulence of the pathogen. In addition, the region should be transcriptionally active to allow proper expression of the cassette.

We have observed in several studies where *Fg* genes functions were characterised that integration of expression cassettes occurred randomly (7, 9, 12). Integrations for *Fg* gene complementation and protein localisation studies are usually done by ectopic integration. The identification of a suitable locus for TSI in *Fg* would avert the potential risks of ectopic integration. Here, we identified a 3.2 kb intergenic region in a highly conserved region on chromosome 1 suitable for TSI. We named this intergenic region the TSI locus 1. We developed an efficient cloning vector system based on the Golden Gate method to clone different expression cassettes for use in combination with TSI locus 1. We present evidence that integrations in the TSI locus 1 affects neither fungal virulence nor fungal growth under different stress conditions. Integrations at the TSI locus 1 successfully resulted in the expression of different gene fusions. In addition, the activities of the trichodiene synthase promoter and effector promoters were not altered by integration into the TSI locus 1. We have developed a bespoke bioinformatic pipeline to analyse the existence of deletions, ectopic integrations and tandem insertions of the cassette that may occur during the transformation process. Finally, we established a protocol to study protein secretion in wheat coleoptiles using confocal microscopy and the TSI locus 1 for stable expression of different gene fusions. In summary, the TSI locus 1 can be used in *Fg* and potentially other cereal infecting Fusarium species for diverse studies including promoter activity analysis, secretion, protein localisation studies and gene complementation.

## Methods

### Strains, media and culture

*Fg* wild strain PH-1 (13) was used for all the transformation events while for complementation analysis the PH1*-*Δosp24-1 mutant was used. Fungal strains were maintained on SNA (synthetic nutrient poor agar) plates. For growth and sporulation of the strains transformed into TSI locus 1 or *osp24* locus, the SNA plates also contained either 75 µg/ml of geneticin (G148, Sigma-Aldrich) or 75 µg/ml of hygromycin B (Calbiochem), respectively as selection agent. Plates were kept under constant illumination (UV and white light) at room temperature (RT). Conidia production on SNA plate was induced by adding 4 ml of TB3 media to 7 days-old mycelia (14). Spores were collected after 24 hours in sterile water and stored at −80 L as described before (14). DNA plasmids were amplified using *Escherichia coli* strain DH5L. *E. coli* transformed cells were selected in LB agar media containing ampicillin 100 µg/ml (Melford, UK) or spectinomycin 150 µg/ml (Melford, UK).

Defects in radial growth in the transformant strains compared to PH-1 were evaluated under different stress conditions. Twenty-five ml of half-strength PDA (Potato Dextrose Agar) containing 2 % agar were mixed with different stress inducing agents such as salt stress (1M NaCl), and membrane stresses (100 µg/ml Calcofluor, 50 µg/ml Congo Red, 0,02 % Tergitol or 0,002 % SDS). The agar mixed with a single stress inducer were poured into squares plates (Grenier Bio-One, UK). Serial spore dilutions were prepared for each transformant and PH-1, and 5 µl of each dilution were plated. A half-strength PDA plate without any stress agent was included as the control. Plates were incubated in a dark cabinet at RT for the entire experiment. Photographs were taken 3 days post inoculation (dpi). The experiment was repeated three times.

### Plasmids design and cloning strategies

To build the *Fg* vector, several cloning steps were performed (Additional file 1: Fig.S1A). First, to generate a geneticin resistance cassette, the gpdA promoter (*P_gpdA_*) and the trpC terminator (*T_trpC_*) were synthesised (Epoch Life Science, US) while the geneticin gene was amplified from the pCGEN vector (15). Next, *BsaI* sites to the *P_gpdA_*, *T_trpC_* and geneticin gene were added using primer combinations pGPDApro_F - pGPDApro_R, TtrpC_F - TtrpC_R, and Gene_F1 - Gene_R1, respectively. All primers used in this study are listed in additional file 2: Table 1. To assemble the geneticin cassette (*P_gpdA_*-geneticin-*T_trpC_*), the Golden Gate protocol was used as described before (16). Finally, from the geneticin cassette a PCR product was amplified containing the *P_gpdA_* and a split fragment of the geneticin gene (*P_gpdA_-* geneticin_1-664_) with primers Gene_R XhoI and Gene_F BsaI. For the second phase of the genetic cassette construction, the right border (RB) of the TSI locus 1 was cloned from PH-1 genomic DNA using primers FgRB_R SapI (P4) and FgRB_F XhoI. The PCR products containing the RB border and the *P_gpdA_-*geneticin_1-664_ were digested with *XhoI* (New England Biolabs, UK) and ligated using T4 DNA ligase (New England Biolabs, UK). During the ligation a Golden Gate cloning site between the RB border and the geneticin fragment was created. The ligation product (geneticin_1-664_-*P_gpdA_-*RB) was amplified by PCR with primers Gene_F SapI (P3) and FgRB_R SapI (P4). The spectinomycin cassette and the bacterial origin of replication (SpecR-Ori) were amplified by PCR from the pGreen vector (17) with primers Dest_F SapI and Dest_R SapI. Finally, to assemble the entire *Fg* vector, the PCR products containing the geneticin_1-664_-*P_gpdA_-*RB and the SpecR-Ori were digested with *SapI* (New England Biolabs, UK) and ligated with T4 DNA ligase. *E. coli* was transformed with the product of the restriction-ligation reaction and the correct clone was selected by sequencing the vector. To build the vector pJET-LB-geneticin, the following steps were performed (Additional file 1: Fig.S1B). From the geneticin cassette a PCR product containing a split fragment of the geneticin gene and the terminator (geneticin_128-795_-*T_trpC_*) was amplified with primers TtrpC_F AgeI and Gene_R2 BsaI. Next, the LB border of the TSI locus 1 was amplified by PCR from PH-1 genomic DNA with primers FgLB_F1 and FgLB AgeI_R. The PCR products containing the geneticin_128-795_-*T_trpC_*and LB border were digested using *AgeI* (New England Biolabs) followed by a ligation reaction. The ligation product (LB-*T_trpC_*-geneticin_795-128_) was amplified by PCR using primers FgLB_F XhoI (P1) and Gene_R XhoI (P2). Finally, the PCR product was cloned into the vector pJET (Thermo Fisher Scientific, UK) following the manufacturer’s instructions. Positive clones containing the pJET-LB-geneticin were selected by sequencing. All PCR amplifications were done using Q5® High-Fidelity DNA Polymerase (New England Biolabs, UK) following manufacturer’s instructions.

**Additional file 2. Table 1.** List of primers used in this work.

Promoter regions were cloned from PH-1 genomic DNA. We cloned 1000 bp upstream of the start codon of the trichodiene synthase (Tri5) and FgramPH1_01G11655 (*Fgeffector1*) genes with primer combinations ProTri5_F1 - ProTri5_R2 and ProFgEffector1_F - ProFgEffector1_R, respectively. The Tri5 promoter (*P_tri5_*) possesses an internal *BsaI* site between −221 and −216 bp upstream of the start codon. To mutate the site (T_-219_A), two fragments were amplified from the cloned promoter region using primer combinations ProTri5_F1 - ProTri5_R1 and ProTri5_F2 - ProTri5_R2. Next, Golden Gate protocol was used as described (16) to mutate the site and assemble at the same time the two fragments. The product of the Golden Gate reaction was amplified with primers ProTri5_F1 and ProTri5_R2. The PCR product containing the *P_tri5_* was used for cloning into the *Fg* vector. The TrpC promoter (*P_TrpC_*) was cloned from plasmid pHYG1.4 (18) with primers PtrpC_F and PtprC_R. *P_TrpC_*and *P_gpdA_*, and terminator *T_trpC_* containing *BsaI* sites were cloned into vector pJET following the manufacturer instructions (Thermo Fisher Scientific, UK). To clone the coding sequence of *Fgeffector1*, cDNA from wheat floral tissue infected with PH-1 was used as template with primers FgEffector1_F and FgEffector1_R. Constructs cloned into the *Fg* vector were done using the Golden Gate protocol and the library of modules as described (16).

To study complementation at TSI locus 1, the recently identified secreted virulence protein coded by the *osp24* gene (19) was deleted in PH-1 using the ‘split-marker’ approach (20). To delete the *osp24* gene, two vectors (pMU487 and pMU488) were designed. Vector pMU487 consists of a DNA fragment containing 1000 bp upstream the start codon of *osp24* gene (*P_osp24_*) followed by the partial sequence of the hygromycin B (Hyg) cassette (*P_trpC_*-Hyg_1-761_). Vector pMU488 consists of a fragment of the Hyg cassette (Hyg_296-1027_) followed by 504 bp of the terminator region of *osp24* gene (*T_osp24_*). Both vectors shared a 466 bp overlapping region of the Hyg cassette to facilitate recombination. To construct these vectors, *P_trpC_*-Hyg_1-761_ and Hyg_296-1027_ fragments were PCR amplified from pHyg1.4 vector (18) using primer combinations O3 - O4 and O5 - O6, respectively. *P_osp24_* and T*_osp24_* sequences were amplified from PH-1 genomic DNA using primer combination O1 - O2 and O7 - O8, respectively. Finally, Gibson assembly was used to fuse the PCR products (*P_osp24_* with *P_trpC_*-Hyg_1-761_ and Hyg_296-1027_ with T*_osp24_*) and ligated them into the pGEM-T Easy vector (Promega UK Ltd). Selection of the deleted strain (PH1-Δosp24-1) was done using the following primer combinations O9 - O10, O11 - O12 and O13 - O14.

The PH1-Δosp24-1 mutant was complemented at the TSI locus 1 with the *osp24* gene containing its native promoter and terminator. A DNA fragment containing the *P_osp24_,* the *osp24* coding sequence and the *T_osp24_* (*P_osp24_-osp24-T_osp24_)* was amplified from genomic PH-1. Two primer combinations: O15 - O16 and O17 - O18 were used to amplify the fragment due to the presence of an internal *BsaI* site in the promoter region (T_-313_A). The site was mutated using the same strategy as for the *Tri5* promoter. Finally, the PCR product containing the *P_osp24_-osp24-T_osp24_* was cloned into the *Fg* vector. All the plasmids developed in this work (Table 2) will be available from Addgene (Massachusetts, USA) once the manuscript is accepted for publication.

**Table 2.** List of vectors developed in this work.

| Name | Purpose | Bacterial resistance |
| --- | --- | --- |
| Fg vector | Destination vector for cloning and transformation into TSI locus 1 | Spectinomycin |
| pJET-LB-geneticin | Vector to amplify the LB-geneticin fragment for transformation into TSI locus 1 | Ampicillin |
| Fg-$P_{trpC}$-mCherry-$T_{trpC}$ | Constitutive expression of a non-secreted version of mCherry | Spectinomycin |
| Fg-$P_{trpC}$-SP<sub>OSP24</sub>-mCherry-$T_{trpC}$ | Constitutive expression of a secreted version of mCherry | Spectinomycin |
| Fg-$P_{Tri5}$-GFP-$T_{trpC}$ | GFP expression controlled by the *Tri5* promoter | Spectinomycin |
| Fg-$P_{FgEffector1}$-FgEffector1-GFP-$T_{trpC}$ | Fgeffector1-GFP expression controlled by the Fgeffector1 promoter | Spectinomycin |
| Fg-$P_{trpC}$-GFP-$T_{trpC}$ | Constitutive expression of a non-secreted version of GFP | Spectinomycin |
| Fg-$P_{osp24}$-*osp24*-$T_{osp24}$ | *osp24* expression under the *osp24* promoter | Spectinomycin |
| pMU487 | Vector to amplify the $P_{osp24}$-$P_{trpC}$-HyG<sub>1-761</sub> fragment for *osp24* gene mutation | Ampicillin |
| pMU488 | Vector to amplify the Hyg<sub>296-1027</sub>-$T_{osp24}$ fragment for *osp24* gene mutation | Ampicillin |
| pJET-$P_{trpC}$ | Vector containing the *TrpC* promoter for Golden gate cloning | Ampicillin |
| pJET-$P_{gpdA}$ | Vector containing the *gpdA* promoter for Golden gate cloning | Ampicillin |
| pJET-$T_{trpC}$ | Vector containing the *TrpC* terminator for Golden gate cloning | Ampicillin |

### Fungal transformation

Integrations into the TSI locus 1 were done following an adaptation of the ‘split-marker’ approach previously described (21). Constructs cloned in the *Fg* vector were amplified by PCR using primer combination P3 and P4. The region containing the LB and a fragment of the geneticin cassette was amplified by PCR from the vector pJET-LB-geneticin using primer combination P1 and P2. For the *osp24* gene deletion, LB and RB fragments were amplified using U874 - U768 and U769 - U868 primers from pMU487 and pMU488, respectively. PCR products were amplified using HotStar TAQ polymerase (Qiagen) following the manufacturer’s instructions. PCR products were adjusted to a concentration of 2 µg/µl. A 5 µl aliquot of each PCR product amplified from the *Fg* vector was mixed with 5 µl of the product amplified from the vector pJET-LB-geneticin. The mixture of PCR products was used to transform 1×10^8^ protoplasts derived from fungal conidia following a previously described protocol (22). Transformants were selected in regeneration media (0.7% agarose, 0.2% Yeast Extract, 0.2% Casein-Hydrolysate (N-Z-Amine A), 0.8M sucrose) containing 75 ug/ml of geneticin. Two days after transformation, six well-spaced transformants were selected and transferred to a 6-well plate containing SNA agar media with 75 µg/ml of geneticin. Aerial hyphal fragments (minus agar) were collected from each transformant, and DNA was extracted using an alkaline-heat DNA extraction protocol. Briefly, hyphae were resuspended in 100 µl of a 50 mM NaOH solution and heated at 95 L for 15 min. Next, 11µl of 1M Tris-HCL (pH 7) was added to the mixture and centrifuged to precipitate hyphae debris. Then, a 1 µl aliquot of the suspension was used to validate the transformants by PCR with four primer combinations. Primer combinations P5 - P6, P7 - P8 and P9 - P10 were used to confirm insertion of the expression cassette into the TSI locus 1. Whereas primer combination P11 and P12 was used to test for homozygosity of the transformants. In the case of the complemented PH-1-Δosp24 strain, the following primer combinations P5 - P6, P7 - P8, P11 - P12, O9 - P10 and O9 - O10 were used to test correct insertion of the osp24 cassette into the TSI locus 1.

### DNA sequence alignments and whole-genome sequence analysis of transformed strains

Multiple DNA sequence alignments of the LB and RB from PH-1 and various *F. graminearum* isolates and other Fusarium species collected from different global locations were done using Clustal Omega tool (23). For whole-genome sequence analysis. Spores of candidate transformants were inoculated in 200 ml liquid YPD complete medium and grown with agitation (180 rpm) for 2 days at 28 L. One gram of fungal biomass was harvested by filtration. DNA was extracted using the Nucleon PhytoPure DNA extraction kit (Cytiva, UK) following manufacturer instructions. Subsequently, the samples were sent to Novogene (Cambridge, UK) for Illumina sequencing. Sequencing was performed using 150-bp paired-end reads, generating 2 G raw data per sample, with PCR-free library preparation. The wildtype strain PH-1 was also included in the sequencing analysis as a control. To identify the genomic region where the expression cassette was inserted during transformation the following steps were taken. First, a quality control of the reads was assessed by FastQC (24). Then, adapters and low-quality reads were removed using Trimmomatic (25) with the following trimming steps: ILLUMINACLIP, SLIDINGWINDOW and MINLEN. Reads belonging to each strain were aligned to the reference genome of PH-1 (YL1 version, NCBI GenBank number: PRJNA782099) (26) using HISAT2 (27). Finally, the read depth at each base of the genome was computed using the option genome coverage from BEDTools (28). Reads aligned to the reference genome were visualised using Integrative Genomic Viewer (IGV) (29). Mapping statistics were calculated for each sequenced strain using Qualimap (30). To identify deletions in the transformant strains, bases with a read depth ≤ 1 were kept using a filter tool from the Galaxy platform (https://usegalaxy.eu/). The filter tool allows to restrict dataset using simple conditional statements. Bases without coverage may indicate chromosomic regions where deletions or insertions of the expression cassette had occurred. To identify tandem insertions of the cassette in the TSI locus 1. The PH-1 genome was edited by inserting in the TSI locus 1 the expression cassette either from the non-secreted version of mCherry (mCherry) or the secreted version of mCherry (SP-mCherry). Then, reads from each strain were aligned against the respective genome sequence. Read depth for each base was calculated as described before. The genomic regions containing the different sections of the expression cassette were filtered and for each section an average read depth value was calculated. Ratio values between the average read depth of each section and the average read depth from two genes (FGRAMPH1_01G06815 and FGRAMPH1_01G06817) flanking the TSI locus 1 were calculated. The ratio values give an approximation of the number of inserted copies of the cassette in the TSI locus 1. Finally, contigs with evidence of tandem insertion in the mCherry strains were identified. Unmapped reads after alignment with PH-1 were extracted and a de novo assemble approach was performed using SPAdes with –isolate option (31). Contigs were blasted against the cassettes of the mCherry and SP-mCherry strains. Contigs containing truncated sequences from the expression cassettes were selected for further analysis. All the tools used to analyse the sequencing data are available at usegalaxy.eu (32). The raw sequencing data was deposited in the European Nucleotide Archive (ENA) and are accessible through series accession number: PRJEB64490.

### Growth of strains to test reporter gene expression

To explore reporter gene expression during *in vitro* growth, liquid cultures were used. These were started by mixing 1 ml of sterile distilled water containing 10^6^ conidia/ml with 9 ml of TB3 media in 50 ml falcon tubes. Cultures were grown by agitation (100 rpm) at 20 L for 16 hours under dark conditions. Then, fluorescence emission from germinated spores was evaluated by confocal microscopy.

### Wheat floral spikes and coleoptiles infection assays

Plants of the susceptible wheat cv. Bobwhite or Apogee were grown in a growth chamber as previously described (33). Wheat plants at the flowering stage were selected for inoculation. We followed the point inoculation protocol (34). Briefly, wheat spikes at anthesis were inoculated with a 5 µl water solution containing 5×10^5^ conidia/ml. Two spikelets per wheat spike were inoculated and five plants per strain. The control plants were inoculated with water only droplets. Point inoculations were done at the 9^th^ and 10^th^ spikelets counted up from the bottom. All inoculated plants were randomised both during the 48 hours high humidity incubation and again after placing on the controlled growth room shelf. The growth room conditions were 23L/18L (day/night), 60% humidity and 16 hours photoperiod (180 μmol m^-2^ s^-1^light intensity). At 3 days post infection, spikelets showing symptoms of Fusarium infection were selected for confocal analysis. To test whether insertion in the TSI locus 1 affected the infection process, we inoculated the wheat plants with either the wild type PH-1 or the PH-1 strain expressing the non-secreted version of mCherry integrated at the TSI locus 1. A minimum of eight plants per strain were inoculated at 12 days post inoculation (dpi), visibly diseased spikelets were counted below the point of inoculation. The experiment was repeated twice.

Wheat coleoptile infection assay was performed following an adaptation of a previously described protocol (35). Wheat seeds of the cv. Bobwhite were vernalised in the dark for 48 hours. Small pieces of cotton wool soaked with sterile water were placed in each well of a 24-well tissue culture plate (VWR International, US). One wheat seed was placed with the crease facing downwards into each well. The plate was placed inside a humidity chamber for 3 days to allow germination and coleoptile elongation. Then, between 2 to 3 mm of the tip of each coleoptile was removed with scissors. A 10 µl clear plastic pipette tip (Starlab, UK) was cut down to 12 mm above the base and a 12 x 15 mm piece of Whatman 1 filter paper (Camlab, UK) was rolled and inserted inside each tip. Each plastic tip was soaked in a solution containing either 5×10^5^ conidia/ml sterile distilled water or just sterile distilled water. An individual pipette tips was placed over the top of each cut coleoptile. This step was found to be necessary to keep high humidity conditions during incubation to ensure conidia germination and successfully infection. In addition, the Whatman paper must be kept throughout in close contact with the coleoptile to allow transfer of the conidia to the host tissue. The prepared plates were then returned to the humidity chamber and incubated in the dark for another 48 hours. After incubation, the plastic tips were removed from the inoculated coleoptiles. Coleoptiles showing visible symptoms of infection were selected for evaluation under the confocal microscopy. Six coleoptiles per strain were infected. The growth room conditions throughout the experiments were the same as the described for wheat spike infections. The experiment was repeated twice.

### Confocal microscopy

Confocal microscopy was used to explore the expression of fluorescent reporter proteins such as mCherry and GFP in liquid cultures and/or in tissue samples taken from inoculated wheat plants, both floral and coleoptile. Strains expressing GFP under different promoters were inoculated on wheat spikes. At three dpi, lemma tissue was isolated using a scalpel blade from spikelets displaying typical symptoms of *Fg* infection. Lemmas were mounted in sterile water into glass slides and GFP emission was observed under confocal microscopy. Protein secretion in wheat coleoptiles were performed with strains expressing either a non-secreted or a secreted version of mCherry. After 2 dpi, the infected epidermis layer was removed from the coleoptile surface in the vicinity of the visible lesion using a surgical blade and mounted onto a glass slide with sterile water. Hyphae tips displaying fluorescence signal were evaluated by confocal microscopy. Germinated spores in liquid media were mounted in TB3 liquid media into glass slides. Fluorescence emission from the hyphae was evaluated under the confocal. The wild type PH-1 strain was included in all the evaluations to set the confocal conditions. Fluorescence emission was observed using a Stellaris 8 Falcon confocal microscope (Leica, UK). Excitation/emission wavelengths were 561 nm / 590-640 nm and 489 nm / 500-530 nm for mCherry and GFP, respectively. Laser intensity was set between 5 and 10 % in counting operating mode for both fluorescence signals. Images were analysed using ImageJ and LAS X v3.7 software from Leica. Liquid culture evaluation and wheat spike infections were repeated twice whilst coleoptile infections were done three times. A minimum of 3 tissue samples / treatment / construct was examined by confocal microscope in each independent experiment.

## Results

### Identification of a conserved micro-region in chromosome 1 suitable for target site integration (TSI)

To identify new virulence factors in *Fusarium graminearum* (*Fg*). Beacham and collaborators developed a bespoke bioinformatic approach that allowed the identification of a micro-region in chromosome 1 enriched in homologues of known virulence genes from multiple cereal and non-cereal disease-causing fungal species (36). The genes were manually curated over a 10-year period into the publicly available pathogen-host interactions database (PHI-base) (36–38). The micro-region spanned the region from FGRAMPH1_01G06783 to FGRAMPH1_01G06811 and contains a total of 15 genes. This micro-region contained homologues of 5 already well characterised fungal virulence genes. In addition, the micro-region was described as residing in a region of low recombination and highly conserved among different Fusarium species such as *F. verticillioides*, *F. oxysporum* and *F. solani* (13, 36). The low recombination rates in the region, the high abundance of virulence gene homologues as well as the high degree of conservation, i.e. make this area a potentially suitable region for target site integration. Just to the 3’ of this micro-region, there is an unusually wide intergenic region of 3274 bp between the predicted genes FGRAMPH1_01G06815 and FGRAMPH1_01G06817. The insertion of an expression cassette in this intergenic region was predicted not to affect the function of the neighbouring genes (Fig. 1A). To permit homologous recombination between the intergenic region and an expression cassette, we decided to use the ‘split-marker’ approach (21). This approach has been used for gene deletion and complementation in *Fg* (34, 39, 40). The approach consists in transforming the fungus with two overlapping DNA fragments. Each fragment is flanked by a ∼1 kb sequence with homology to the target locus. The flanked sequences are defined as left and right borders (LB and RB, respectively). Insertion of the expression cassette in the target locus occurs by a triple homologous recombination event. Two events occur between the LB and RB with their respective homologous sequences in the target locus. A third event occurs between the two overlapping DNA fragments (Fig. 1A). We built a vector system to adapt this methodology for target site integration into the 3274 bp intergenic region, now referred to as target site integration locus 1 (TSI locus 1). One vector called pJET-LB-geneticin contains an 816 bp DNA fragment of the *Fg* genome as the LB followed by a resistance cassette. The resistance cassette has the *Aspergillus nidulans* trpC terminator (*T_trpC_*) and a 667 bp split fragment of the geneticin gene as the selection marker (Fig. 1B). The second vector called the *Fg* vector contains a 664 bp split fragment of the geneticin gene where 536 bp overlaps with the sequence of the geneticin fragment in pJET-LB-geneticin. The 664 bp geneticin fragment is followed by the *A. nidulans* constitutive gpdA promoter (*P_gpdA_*) and an 848 bp DNA fragment of the *Fg* genome as the RB (Fig. 1B). Between the promoter *P_gpdA_* and the RB, there is a cloning site (CS) adapted to the Golden Gate approach (41) where the type IIS enzyme *BsaI* cuts twice the vector creating two single-stranded overhangs with 4 bp each. The overhang sequences were defined according to those previously described (16) and thus the modular library developed by the authors can be also used for cloning in the *Fg* vector (Fig. 1B). To evaluate if this vector system could or could not be used for transformation of other *Fg* isolates and Fusarium related species, we aligned the LB and RB sequences from PH-1 with the corresponding genomic regions and sequences from isolates of *Fg, F. culmorum* and *F. pseudograminearum* collected worldwide. Both borders are conserved among different *Fg* isolates indicating that the vector system can be used to transform strains from different origins. In addition, the degree of conservation in *F. culmorum* and *F. pseudograminearum* is also high (Additional file 1: Fig.S1C). In both species however, a small insertion of 116 bp in the 5’ of the LB sequence was identified, but we consider that there is enough homology to adapt this strategy for transformation of both species.

**Figure 1.**
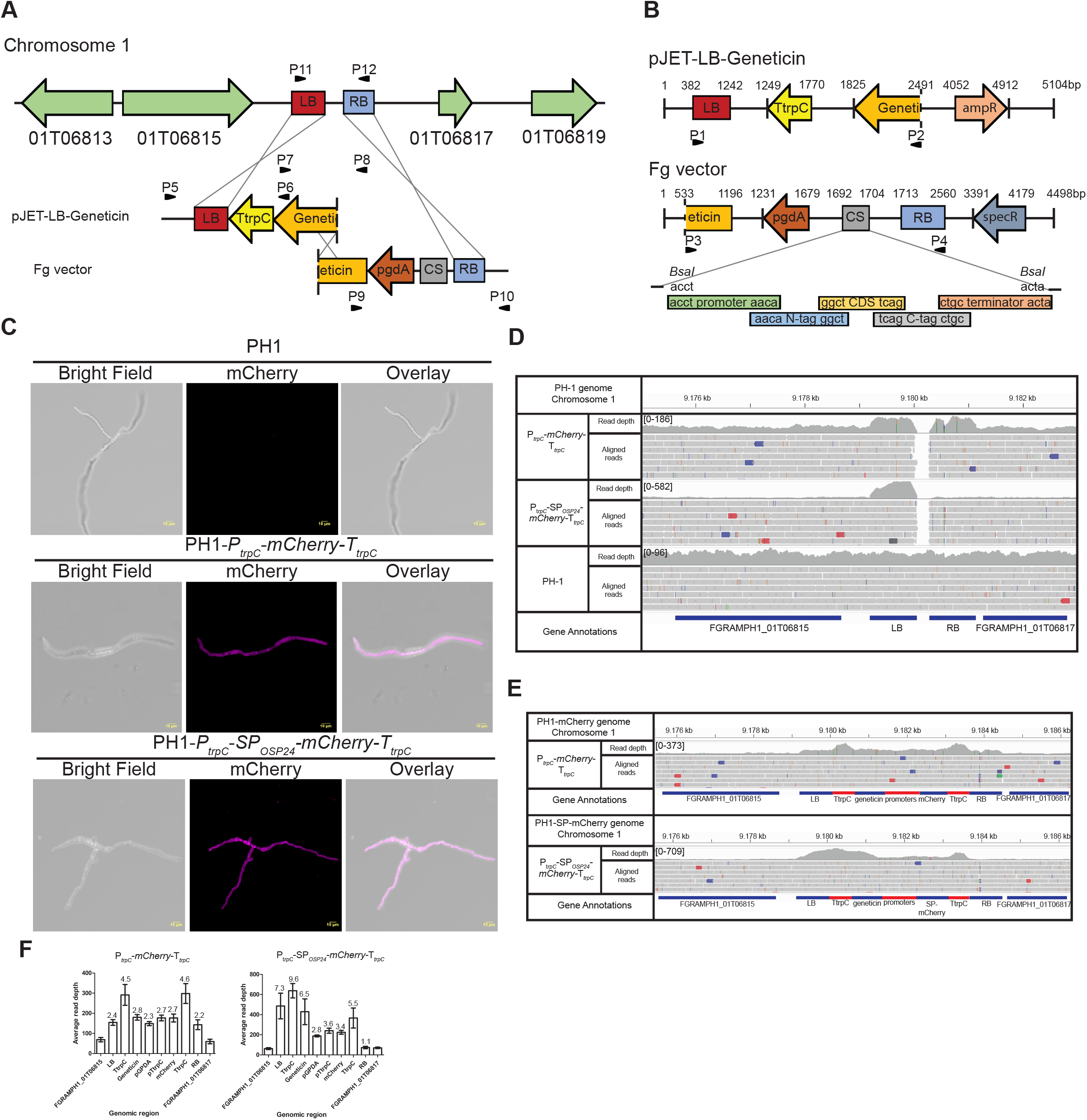
Schematic representation of the locus for TSI and confocal analysis of *F. graminearum* transformants. **A)** The TSI locus 1 is located in a micro-region within chromosome 1 between the genes designated FGRAMPH1_01G06815 and FGRAMPH1_01G06817. Inside this micro region, there is an intergenic region of 3.2 kb where the insertion of the expression cassette can occur. To allow integration of the expression cassette into the TSI locus 1, a vector system was developed based on the split-marker technique. Three recombination events allow insertion of the cassette into the TSI locus 1. Primer combinations P5-P6, P7-P8, P9-P10 and P11-P12 are used to confirm correct cassette integration. **B)** A vector system based on the Golden Gate approach was developed to allow cassette integration into the TSI locus 1. PCR fragments amplified by primer combinations P1-P2 and P3-P4 are used for fungal transformation. **C)** Confocal images of strains confirming expression of a non-secreted version of mCherry (*P_trpC_-mCherry-T_trpC_*) and a secreted version (*P_trpC_-SP_OSP24_-mCherry-T_trpC_*). Fluorescence emissions were detected 16 hours later. Wild type strain PH-1 was used as the control to set the confocal conditions. **D**) IGV screenshot showing the reads aligned to the LB and RB of the TSI locus 1 in wild the type strain PH-1 and the transformed strains. Numbers in brackets indicate the range of read depth coverage per bp. **E**) IGV screenshot displaying reads aligned to the expression cassettes from the transformed strains and the genes flanking the TSI locus 1. **F**) Bar graphs represent the average read depth values for each section of the expression cassette and for the two genes (FGRAMPH1_01G06815 and FRGAMPH1_01G06817) flanking the TSI locus 1 for both transformants. Values above the bars are the ratio value calculated as the average read depth value of each section divided by the average read depth value from both flanking genes. Error bars represent standard deviation.

### Transformation into the TSI locus 1

To test if the TSI locus 1 is a suitable region for transformation and gene expression, we built two constructs in the *Fg* vector. A non-secreted version of mCherry (mCherry) under the control of the constitutive promoter of *A. nidulans* trpC (*P_trpC_*) flanked at the 3’ end by the terminator sequence *T_trpC_* (*P_trpC_-mCherry-T_trpC_*). A second construct consisted of a secreted version of mCherry (SP-mCherry) where we used the 69 bp secretion signal from the orphan secreted protein 24 (*SP_osp24_*) (19). The expression of mCherry in this second construct is also controlled by *P_trpC_*and has the terminator sequence *T_trpC_* (*P_trpC_-SP_osp4_-mCherry-T_trpC_*).

Each cassette was amplified by PCR using the overlapping DNA fragments from pJET-LB-geneticin and the constructs cloned into *Fg* vector using primer combination P1 - P2 and P3 - P4, respectively (Fig. 1B). The transformation of the wild type PH-1 strain resulted in ∼20 geneticin resistant colonies from two 25 ml square plates and six colonies were selected from each transformation event. To validate that the transformations were successful, colonies were evaluated by diagnostic PCR. Primer combinations P5 - P6 and P9 - P10 were used to confirm that insertion had occurred in the TSI locus 1 (Fig. 1A and Additional file 3: Fig. S3A). The primer combination P7 - P8 was used to confirm that the recombination event between the two overlapping DNA fragments was correct (Fig. 1A and Additional file 3: Fig. S3A). *Fg* protoplasts might contain more than one nucleus during the transformation and thus not all the nuclei can be transformed during the transformation step. To check that the colonies selected are homokaryotic for the expression cassette, PCR analyses were done on a region between the LB and RB using the primer combination P11 - P12. The lack of an 868 bp band belonging to untransformed nuclei showed that all the colonies were homokaryotic (Fig. 1A and Additional file 3: Fig. S3A). To test for the expression of mCherry, we evaluated mCherry fluorescence emission by confocal microscopy in spores germinated in liquid TB3 media. The colonies selected for each transformation event displayed fluorescence emission (Fig. 1C).

To verify target site integration of the expression cassette in the TSI locus 1 and assess whether deletions might have occurred during transformation, the genomes of the mCherry and SP-mCherry strains were sequenced. The sequencing reads from each strain were aligned to the reference genome of the wild type strain PH-1. Around 99% of the reads from each strain were mapped to the wild type genome to a mean coverage value of 54-65x for each chromosome (Table 3). For both strains, reads covered the whole genome of PH-1 except for 228 bases between the LB and RB of the TSI locus 1 (Fig. 1D), indicating insertion of the cassette only in the TSI locus 1. Other genomic regions with low read coverage were also detected in the transformant strains (Additional file 4: Fig. S4A). However, these regions were also presence in the aligned sequences obtained from the wild type PH-1 strain. Therefore, we can conclude that there is no evidence of ectopic integration or deletions in both transformant. To estimate the number of copies of the cassette inserted in the TSI locus 1. The reads from each strain were aligned to the PH-1 genome containing the sequence of the respective expression cassette inserted in the TSI locus 1. The expression cassettes from both strains were fully covered by reads indicating that both cassettes are complete (Fig. 1E). This evidence agrees with the mCherry expression observed for both strains (Fig. 1C). However, it was observed that the number of reads aligned to the expression cassettes was higher than the number of reads aligned to two neighbouring genes (FGRAMPH1_01G06815 and FGRAMPH1_01G06817) flanking the TSI locus 1 (Fig. 1F). In the case of the mCherry strain, the ratio values between the average read depth for each section of the cassette and the average read depth for FGRAMPH1_01G06815 and FGRAMPH1_01G06817 genes were 2 to 3 times higher (Fig. 1F). This evidence indicates that there are between 2 to 3 copies of the cassette. In the case of the SP-mCherry strain, multiple insertions of the cassette were also observed (Fig. 1E). Ratio values for the regions containing the mCherry gene and the promoters (*P_gpdA_* and *P_trpC_*) were 3 times higher indicating at least 3 copies of the cassette (Fig. 1F). In addition, the ratio for the genomic regions containing the LB, the geneticin gene and one of the terminators were around 7 times higher in comparison to FGRAMPH1_01G06815 and FGRAMPH1_01G06817 genes (Fig. 1F). This evidence indicated that the integration event not only contains copies of the full cassette, but also fragments containing the LB region, the geneticin gene and the TrpC terminator. For both strains the average read depth for the TtrpC terminators was always double the average read depth values from other sections of the cassette (Fig. 1F). The higher values are due to reads that can be aligned to both terminator sequences. Finally, coloured reads were observed in the TSI locus 1 region and in chromosomic regions with low read coverages such as telomeres (Fig. 1D and 1E; Additional file 4 and 5: Fig. S4A, S5C and S5D). Coloured reads displayed in the IGV screenshots might indicate potential structural variants such deletions (red), insertions (blue), inversion (gray-blue) and duplications or translocations (green) (29). In the case of the TSI locus 1 region, coloured reads are only a few and randomly distributed. Therefore, it is assumed that are not true variants. In the case of regions with low coverage, coloured reads were present not only in the transformed strains but also in the wild type (Additional file 4 and 5: Fig. S4A and S5D). Hence, coloured reads could be the consequence of poor coverage or differences with the reference genome used to align the reads.

**Table 3.** Mapping statistics for the strains sequenced.

| Transformant strains | Reads after trimming | Reads mapped (%) | Chromosome mean coverage (mean ± sd) | | | |
| --- | --- | --- | --- | --- | --- | --- |
|  |  |  | Chr1 | Chr2 | Chr3 | Chr4 |
| PH1 | 18.205.628 | 99.73% | 66.72x±11.3 | 65.70x±11.1 | 66.07x±11.1 | 64.34x±16.6 |
| *P<sub>trpC</sub>-mCherry-T<sub>trpC</sub>* | 17.665.176 | 99.70% | 64.18x±11.2 | 63.57x±11.1 | 63.77x±11.1 | 65.33x±14.7 |
| *P<sub>trpC</sub>-SP<sub>OSP24</sub>-mCherry-T<sub>trpC</sub>* | 15.251.700 | 99.70% | 54.97x±10.5 | 54.46x±9.9 | 54.69x±9.9 | 53.81x±13.4 |
| PH1-Δosp24-1 | 13.524.682 | 99.71% | 48.37x±9.1 | 48.03x±9.1 | 48.23x±9.1 | 47.04x±12.1 |

To test for the existence of tandem insertion of the cassette in the mCherry and SP-mCherry strains, unmapped reads were recovered after alignment with the PH1 genome. A de novo assembly approach was performed using the unmapped reads to identify contigs with evidence of tandem insertion. In the case of the mCherry strain, a contig of 263 bp (contig_42) containing truncated sequences of the RB (RB_727-852_) and LB (LB_1-117_) was identified (Additional file 4: Fig. S4B). This contig indicates a head-to-tail tandem insertion of the cassette. In the SP-mCherry strain three contigs were identified. A contig of 366 bp (Contig_11) containing truncated sequences of the LB (LB_248-130_) and RB (RB_131-383_) indicates the existence of a head-to-tail tandem insertion of two copies of the cassette (Additional file 4: Fig. S4B). A second contig of 396 bp (Contig_9) was identified containing truncated sequences of the LB (LB_723-861_ and LB_1-61_) and *TrpC* terminator (*TrpC*_1-180_). Finally, a third contig of 200 bp (Contig_15) containing truncated sequences of the geneticin gene (Gen_204-127_) and the LB (LB_1-118_) was also identified (Additional file 4: Fig. S4B). This evidence indicates the existence of fragments containing truncated sequences from the LB, geneticin gene and TrpC terminator and thus explain the higher number of read mapped onto that region of the cassette.

To test if integrations in the TSI locus 1 could affect fungal infection and disease symptom causing ability, a floral point inoculation test was done. Wheat spikes of the susceptible cv Bobwhite were inoculated with either wild type untransformed PH-1 and the strain expressing the non-secreted version of mCherry. There were no differences in the number of infected spikelets between PH-1 and the mCherry strain indicating that integration in the TSI locus 1 does not affect the virulence of the fungus (Additional file 3: Fig. S3B). We also test if integrations in the TSI locus 1 affected the fungal morphology or growth rate when the fungus was grown *in vitro* under different stress conditions. PH-1 and the transformants were grown in PDA plates containing either salt or membrane stresses. The transformants showed a similar morphology and growth rate as PH-1 for all the conditions tested (Additional file 3: Fig. 3C). Hence, insertions in the TSI locus 1 does not affect the fungal growth under the conditions tested.

### Validation of protein secretion in *F. graminearum*

The yeast secretion trap assay is a commonly used method to validate protein secretion (42). However, in plant pathogens such as the fungus *U. maydis* and *M. oryzae*, secretion studies are performed in the native system. A protein predicted to be secreted is fused to a fluorescence reporter and the fungus is transformed with the recombinant protein. If the protein is secreted, an accumulation of the fluorescence signal can be observed in the periphery of the infected hyphae whilst a non-secreted protein will only accumulate inside the hyphae (43–45). We evaluate whether the *Fg* strains expressing the secreted and non-secreted versions of mCherry show similar or dissimilar distribution pattern. We infected wheat coleoptiles with the non-secreted version of mCherry (*P_trpC_-mCherry-T_trpC_*) and the secreted version (*P_trpC_-SP_osp24_-mCherry-T_trpC_*). The strain expressing the non-secreted version displayed accumulation of the mCherry fluorescence signal inside the hyphae. However, the strain expressing the secreted version displayed accumulation of the fluorescence signal in the periphery of the hyphae mainly localised towards the tips (Fig. 2). These results indicate that secretion studies can also be performed in *Fg* using cassettes expressed from the TSI 1 locus.

**Figure 2.**
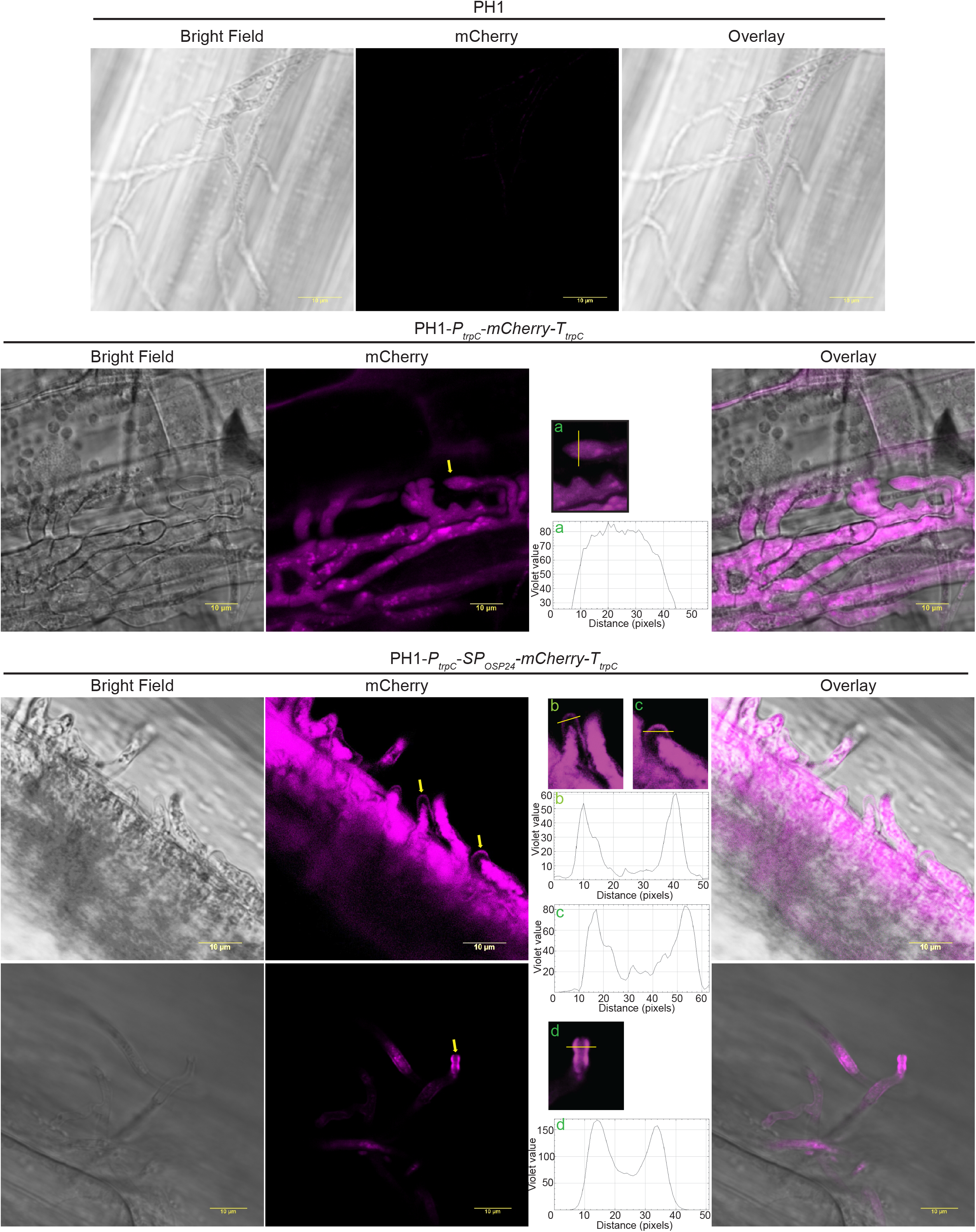
Secretion of mCherry in wheat coleoptiles. Confocal images of wheat coleoptiles infected with the strains expressing either the non-secreted version of mCherry (*P_trpC_-mCherry-T_trpC_*) or the secreted version (*P_trpC_-SP_osp24_-mCherry-T_trpC_*). The strain expressing the non-secreted version displayed accumulation of the mCherry fluorescence signal solely inside the hyphae. Whereas the strain expressing the secreted version displayed accumulation of the fluorescence signal in the periphery mainly localised towards the tips. Graphs a, b, c and d indicate mCherry signal intensity determined along the diameter of the hyphae, taken at the positions indicated by yellow lines in each image enlarged and labelled with the same letter. The yellow arrows point to the selected hyphae used for this analysis. Wild type strain PH-1 was used to set confocal conditions. Confocal images were taken 2 dpi.

### Complementation of PH1**Δ**osp24 by insertion into the TSI locus 1 restores full virulence

Previous studies have shown that o*sp24* (FGRAMPH1_01G15939) is required for successful infection and disease formation within of wheat floral tissue (19). Therefore, the o*sp24* gene is a good candidate to test if the TSI locus 1 can be used for complementation analysis. The o*sp24* gene was mutated in PH-1 by the split marker approach. Analysis by PCR confirmed that the coding sequence of o*sp24* was replaced by the hygromycin cassette (Additional file 5: Fig S5A). Further, sequencing analysis of the mutant strain (PH1-Δosp24-1) showed a 480 bases gap in the coding sequence of *osp24* (Additional file 5: Fig. S5C). The Hyg cassette was integrated in the o*sp24* locus (Additional file 5: Fig. S5C). In addition, there is no evidence of ectopic integration as genomic regions with low coverage in PH1-Δosp24-1 were also present in PH1 (Additional file 5: Fig. S5D). Finally, the Hyg cassette was inserted as a single copy as the ratio values between the different sections of the cassette (LB, hygromycin cassette, and RB) and the genes (FRGRAMPH1_01G15937 and FGRAMPH1_01G15941) flanking the o*sp24* locus were close to 1 (Additional file 5: Fig. S5C). *In planta* testing of this mutant strain in Apogee spikelets showed reduced virulence compared to the wild type PH-1 strain (Fig. 3A). The infection was limited to the inoculated spikelets as previously reported (19). To test for the full restoration of virulence, PH1-Δosp24-1 mutant strain was complemented by inserting the expression cassette (*P_osp24_-osp24-T_osp24_*) containing the wildtype gene into the TSI locus 1. The complemented strains were selected by PCR to confirm correct insertion of the *P_osp24_-osp24-T_osp24_* cassette into the TSI locus 1 (Additional file 5: Fig. S5B). The complemented strain PH-1-Δosp24-osp24-1 regained the wildtype virulence phenotype (Fig. 3A). Hence the TSI locus 1 can be used for efficient complementation studies.

**Figure 3.**
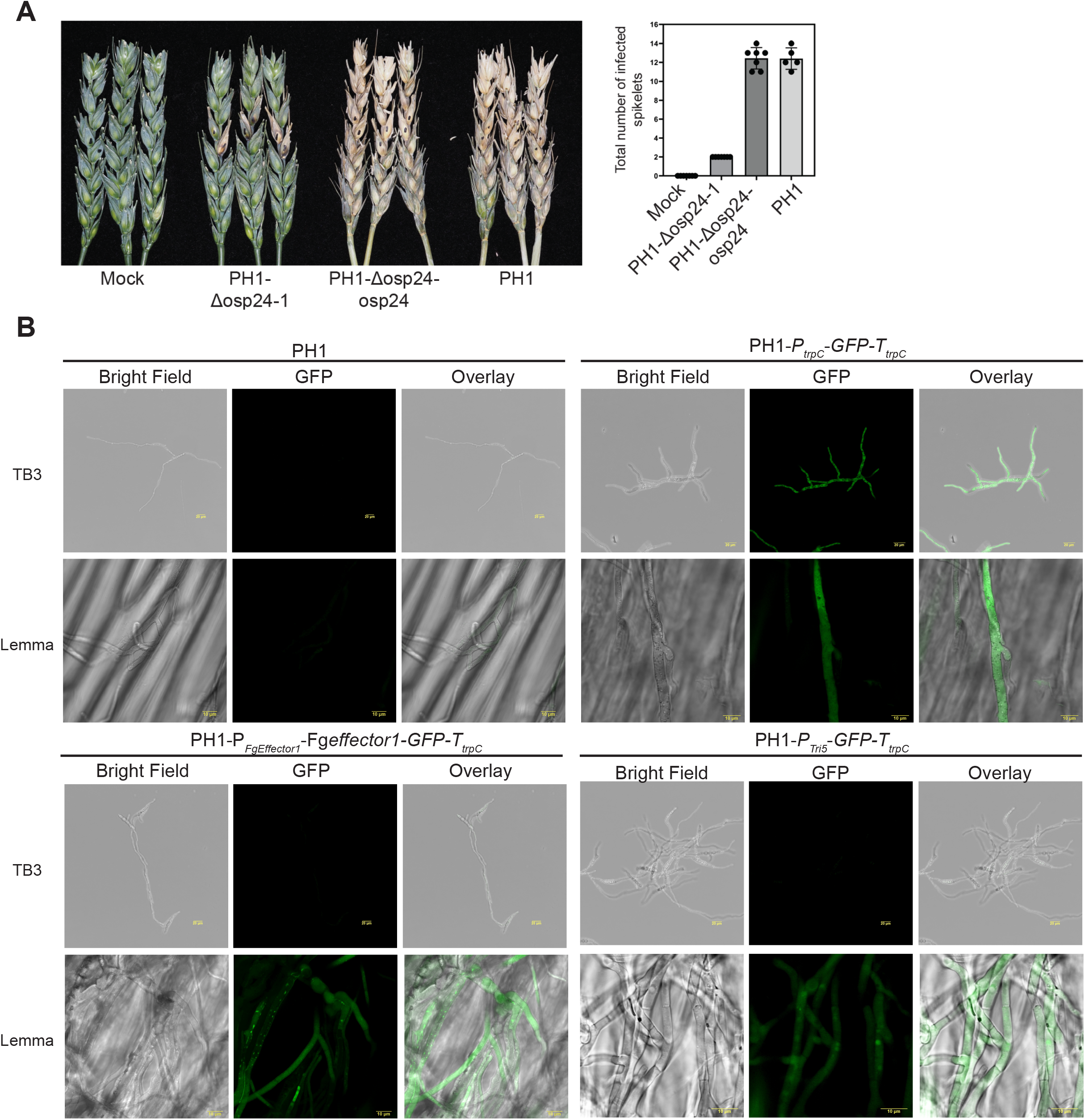
Complementation of PH1-Δosp24-1 and promoter analyses under different conditions. **A)** Complementation of the PH1-Δosp24-1 mutant strain with the *osp24* gene residing within the TSI locus 1 (PH1-Δosp24-osp24-1) restores full virulence. Photographs were taken at 14 dpi. Marked spikelets of each floral spike indicate the inoculation points. Bar graph shows no differences in the number of infected spikelets between PH-1 and PH1-Δosp24-1-*osp24* whilst the mutant strain shows reduced virulence. Visibly diseased spikelets were counted including the point of inoculation after 14 dpi. Mock indicates plants inoculated with water. **B)** Confocal images of strains expressing GFP under the control of different promoters. Strain expressing constitutive GFP (*P_trpC_*-GFP-*T_trpC_*) displayed fluorescence when *F. graminearum* was grown in TB3 liquid media as well as during wheat spike infection (lemma). Expression of GFP under the control of the trichodiene synthase promoter (*P_Tri5_*-*GFP*-*T_trpC_*) or an effector promoter (*P_Fgeffector1_*-*FgEffector1*-*GFP*-*T_trpC_*) only occurs during infection. The PH-1 strain was used as control to set confocal conditions. Images were taken at 3 dpi.

### Virulence specific promoter activity is not altered by insertion into the TSI locus 1

Pathogens possess genes that are exclusively expressed during host infection. *Fg* produces different trichothecene mycotoxins that are required for successful infection of wheat spikes. The *tri5* gene codes for an enzyme that catalyses the first step in the production of all trichothecene mycotoxins (46, 47). The *tri5* gene is highly expressed during infection (48), but only a low expression level is observed in liquid culture unless transferred to specific induction media (49) or by the addition for specific inducers to the cultures for example agmatine (50) or hydrogen peroxide treatments (51). To test if native promoter activity might be affected by insertion in the TSI locus 1, we cloned in the Fg vector, a construct where GFP expression is under the control of the *tri5* promoter (*P_Tri5_*-*GFP*-*T_trpC_*). In addition, we cloned a promoter from a candidate effector gene FgramPH1_01G11655 (*P_Fgeffector1_*) that according to transcriptomic analysis is upregulated during the symptomatic phase of the wheat spike infection (52). A second construct was built where P*_FgEffector1_* controlled the expression of the FgEffector1 gene which was also C-terminally tagged to GFP (*P_Fgeffector1_*-*Fgeffector1*-*GFP*-*T_trpC_*). Finally, as an additional control a construct was generated where GFP was under the control of the *P_trpC_* (*P_trpC_*-GFP-*T_trpC_*). Positive transformants were obtained for the three different constructs (Additional file 3: Fig. S3A) and the transformant strains showed a similar morphology and growth rate as PH-1 for all the nutrient and stress conditions tested (Additional file 3: Fig. S3C). To test promoter specificity, selected strains for the three different constructs were grown in TB3 liquid medium. Only the strain expressing constitutive GFP displayed a fluorescence signal. However, when the same strains were used to infect wheat spikes, all the strains displayed fluorescence, indicating that the activities of the *P_Tri5_* and *P_Fgeffector1_* was not altered by the localisation of either promoter in the TSI locus 1 (Fig. 3B).

## Discussion

*Fg* is an important disease-causing pathogen worldwide that impacts on global food and feed security. The functional characterisation of genes and proteins in this pathogen is required in order to develop novel resistance strategies. Common approaches to study gene/protein functions include gene complementation and protein localisation. These approaches include the integration of expression cassettes into the fungal genome, but frequently these integration events are non-targeted and therefore occur randomly. The identification of a suitable locus for target site integration (TSI) for *Fg* is required to avoid the potential risks of ectopic integration. Another advantage of TSI is that it permits a direct comparison of promoter activities or gene functions under different genetic backgrounds. In this study, the TSI locus 1 located within chromosome 1 was shown to be a suitable region for cassette insertion as we observed good levels of expression for six different cassettes. In addition, we also observed that the *in planta* specific expression of three promoters (P*_osp24_*, P*_tri5_* and P*_Fgeffector1_)* was not affected by insertion in the TSI locus 1. We did not observe phenotypic differences between the transformed strains and the wild type for the different stresses evaluated and during wheat infection suggesting integration of the expression cassette in the TSI locus 1 may not affect the expression of the genes flanking the TSI locus 1. Furthermore, full virulence was restored when the osp24 mutant was complemented with a full copy of the *osp24* wild type gene targeted to TSI locus 1. This locus can be used for the transformation of different *Fg* strains due to the high levels of sequence conservation. In addition, the locus is conserved in two other phytopathogenic Fusaria, namely *F. culmorum* and *F. pseudograminearum* indicating that both species could also be transformed using the same Golden Gate based vector system. In summary, the TSI locus 1 can be used for diverse studies including promoter activity, gene complementation and protein localisation studies.

Multinucleate fungus such as Fg possesses several nuclei per cell and thus it is possible to obtain heterokaryon strains after transformation (15). In addition, during transformation chromosomal rearrangements may occur leading to off-target gene modifications (53). In this study, we did not detect heterokaryotic strains for the different transformation events indicating that the strains were homokaryotic for the expression cassette. In addition, whole genome sequencing analysis showed insertion of the expression cassette only in the TSI locus 1 without any evidence of ectopic integration elsewhere in the genome. However, we have identified multiple tandem insertion of the expression cassette. This was not surprising because the mCherry expressing strains were first selected by scoring the intensity of the fluorescence signal evident during plant infections. Therefore, strains with multiple copies of the expression cassette were selected. It is noteworthy that multiple tandem insertion of the cassette during transformation has been reported in other fungi where the selection marker could be responsible for the concatenation (54–56). In conclusion, the multiple insertion of the cassette observed for the mCherry and SP-mCherry strains are not related with integrations in the TSI locus 1, but it could be a consequence of the prior screening criteria (high fluorescence expression) and/or antibiotic selection during transformation. Southern blot analysis is the gold standard technique used to test for ectopic integration and predict number of copies inserted during transformation (56–58). However, the technique cannot detect structural variants such as deletions occurring elsewhere in the genome during the transformation process. Recently, a whole genome sequencing analysis allowed the detection of a large-scale deletion in Fusarium mutant strains where genes were mutated using CRISPR/Cas9 (59). In contrast, in this study, we were not able to detect deletions in the two mCherry expressing strains or the osp24 deleted strain generated by using the split marker approach. This evidence is consisted with recent findings (54) where the potential mutagenic inheritance of the split marker approach was assessed in the fungus *Cryptococcus neoformans*. The authors sequenced different deletion strains and strains transformed in an apparently neutral locus. No deletions were observed in any of the strains. Only a low but significant point mutation frequency for the deleted strains was found (1.67 point mutations per gene deletion strain) indicating that the split marker approach for gene deletion and target site integration is safe. Finally, by following a bespoke bioinformatic pipeline, we were able to rule out the existence of ectopic integration, verify the insertion of the expression cassette in the TSI locus 1 and predict the number of copies inserted in the TSI locus 1. The identification of ectopic chromosomic regions with deletion or insertion of the expression cassette was possible due to the use of PCR-free libraries that avoid potential bias incurred during library preparation and sufficient sequencing depth (47-66x coverage for the sequenced strains). Even though, we were not able to fully characterize the insertion events for the mCherry and SP-mCherry strains due to the short-read (150 bp) sequencing approach used, we were able to identify contigs containing proof of multiple tandem insertions. Long-read sequencing approaches such as PacBio or Nanopore would enable a better characterization of the insertion events. However, we do not consider relevant a fully characterization of the events as the aim was to obtain transformed lines that express sufficient fluorescent signal to be detected during plant infections. In conclusion, 150 bp paired-end sequencing combined with PCR-free library preparation and 50x read coverage of the genome is a useful approach to assess ectopic integration events as well as to differentiate between single and multi-copy number insertion events. Therefore, the sequencing approach could serve as an alternative to Southern blotting analysis.

Currently, the well-established yeast recombinational cloning approach seems to be one of the preferred options for gene fusions in different *Fg* studies (7–9). The methodology exploits yeast homologous recombination to fuse a gene with different promoters or tags (60). However, this cloning strategy is somewhat tedious because the first cloning step has to be done in yeast. Then, the plasmid is purified, transformed into *E. coli,* amplified to permit the cloned product to be sequenced. We have developed a vector system for integration into the *F*g TSI locus 1 using the Golden Gate cloning approach. The methodology allows the assembly of a gene of interest with different tags and/or promoters in a restriction-ligation reaction without the necessity of using yeast. In addition, the methodology reduces the overall cloning time as allows the use of a library of modules already available. Even though we have not tested the maximum length of the expression cassette that can be inserted into the TSI locus 1. In our hands the protocol was highly efficient for all the constructs tested.

Pathogens secrete many different types of proteins during infections. Candidate secreted proteins are usually identified bioinformatically by the predicted presence of a secreted signal at the N terminus (61). Validation of protein secretion is commonly assessed by using the yeast secretion trap assay. The approach consists of fusing the cDNA of a potential secreted protein to the yeast invertase (suc2) reporter gene lacking its signal peptide. If the protein is secreted, this extracellular targeting allows the growth of a yeast strain defective in suc2 on a sucrose selection media (42). In *Fg*, validation of protein secretion is often done using the yeast secretion trap assay (9, 19, 62). Even though the technique is widely accepted, these heterologous results should ideally be verified with experiments performed in the native system. Secretion studies have been developed for different pathogens such as *U. maydis* and *M. oryzae* (43–45). To explore if this technique could be adapted to *Fg*, we designed two different strains expressing a secreted and a non-secreted version of mCherry. We found that hyphae tips harbouring the secreted mCherry version displayed accumulation of the fluorescence signal in the periphery whilst the non-secreted strain accumulated fluorescence signal inside the hyphae. These comparative results indicate that the technique might be also by exploitable for *Fg*. The technique was performed in wheat coleoptiles; a tissue easy to manipulate and visualise under confocal microscopy. However, not all *Fg* genes may display a similar expression pattern among different host tissues (63). Therefore, the future identification of *Fg* promoters highly induced during coleoptile infection would be required to apply this technique successfully as a semi-high throughput reliable assay for testing *Fg* protein secretion. In addition, a major challenge faced during the setting-up of the technique was the infection strategy *per se* of *Fg*. Both, *U. maydis* and *M. oryzae* infect single cells at earlier time points of infection, allowing the easy identification of hyphae tips and strong fluorescence signal accumulation around the hyphae (64, 65). However, *Fg* grows very fast throughout the infected tissue producing various types of infective hyphae and structures making it difficult to find hyphal tips with strong fluorescence signals. In our studies, the best results were obtained 48 hours post infection in colonised areas where hyphae were growing exclusively intracellularly.

## Conclusion

In this study, we characterised and functionally tested a locus for TSI in *F. graminearum,* the first described for this pathogen. Cassette insertion into the TSI locus 1 does not affect fungal virulence and growth under different stress conditions. We observed good levels of expression for all the expression cassettes tested. In addition, promoter activities were not affected by insertion in the TSI locus 1. The high degree of sequence conservation of the locus would allow the transformation of different *Fg* isolates and potentially some other phytopathogenic Fusarium related species. We developed a vector system for efficient cloning and transformation into the TSI locus 1 and we designed a bespoke bioinformatic pipeline that could be used as an alternative to Southern blotting analysis. Finally, we established a protocol for protein secretion studies using confocal microscopy and tested the suitability of the TSI locus 1 for stable expression of different gene fusions. Hence, the TSI locus 1 and the new modular Fg vector system are versatile tools to study gene/protein functions in *F. graminearum*.

**Additional file 1. Cloning steps to build Fg vector and pJET-LB-Geneticin vectors, and LB and RB sequences alignment. A)** A geneticin resistance cassette was built using the Golden Gate cloning approach followed by PCR to amplify a product containing the *gpdA* promoter and a fragment of the geneticin gene (*P_gpdA_-*geneticin_1-664_). Next, the RB of the TSI locus 1 was amplified from PH-1 genomic DNA. The PCR products from the RB and *P_gpdA_-* geneticin_1-664_ were digested and ligated. During the ligation process a Golden Gate cloning site (CS) was created. A fragment containing a spectinomycin resistance cassette (specR) and a bacterial origin of replication (ori) was amplified by PCR from pGreen vector. Finally, the specR-ori and the geneticin_1-664-_*P_gpdA_*-CS-RB PCR products were digested and ligated to build the Fg vector. **B)** From the geneticin cassette a PCR product was amplified containing a fragment of the geneticin gene fused to the *TtrpC* terminator (*TtrpC*-geneticin _128-795_). The LB of the TSI locus 1 was amplified by PCR from PH-1 genomic DNA. The PCR products containing the LB and the *TtrpC*-geneticin _128-795_ were digested and ligated. The ligation product was amplified by PCR and cloned into pJET. **C)** DNA sequence alignments of the LB and RB from PH-1 and various *F. graminearum* isolates and other Fusarium species collected from different global locations. Asterisks indicate positions which have a conserved base among all the isolates and species. *Fusarium graminearum* (F. gram) isolates: GZ3639 (US, GenBank-accession: AACM00000000); MDC_Fg1 (France, GenBank-accession: UIHA00000000.2); CS3005 (Australia, GenBank-accession: JATU00000000); CML3066 (Brazil, GenBank-accession: LT222053) and Fg-12 (China, GenBank-accession: PRJNA743144). *Fusarium culmorum* (F. cul) UK99 (UK, GenBank-accession: PRJEB12835) and *Fusarium pseudograminearum* (F. pseudo) Fp22-2F (China, GenBank-accession: SRX18242685).

**Additional file 3. Validation of the transformant strains, floral virulence test and stress evaluations. A)** To select transformants where the cassette was correctly inserted into the TSI locus 1, we amplified four different PCR products. Primer combinations P5 - P6 and P9 - P10 were used to verify the insertion event. Primer combination P7 - P8 evaluated whether the recombination event between the two PCR fragments had been successful. Primer combination P11 - P12 tested whether each transformant was homokaryotic for the transgene. Red asterisks indicate the expected PCR size bands. The transformant in lane 5 displayed a slightly higher band size for primer combination P9 - P10 when compared to the other transformants. The higher band in lane 5 could be the product of unequal crossover in the RB region of the cassette. **B)** Wheat spikes inoculated with PH-1 or the transformant strain (*P_trpC_-mCherry-T_trpC_*) did not show differences in the number of infected spikelets showing typical disease symptoms. Photographs were taken at 12 dpi. Bars graph shows no differences in the number of infected spikelets between PH-1 and the transformant strain. **C)** All the transformed strains showed a similar morphology and growth rate as wild type PH-1 for all the conditions tested. Photographs were taken after 3 dpi. Salt stress (NaCl), membrane stresses (Calcofluor, Congo Red, Tergitol, SDS). PDA: potato dextrose agar only.

**Additional file 4. Genomic sequence analysis of the mCherry and SP-mCherry expressing strains. A**) IGV screenshots displaying genomic regions with low read depth coverage. Regions with low read depth were identified not only in the transformant strains but also in PH-1. Low coverage regions were usually identified at the telomeres (Chr1I, Chr1IV, Chr2, Chr3I, Chr3II and Chr4IV), in the 5’UTRs (Chr1II, Chr1III and Chr4I) or 3’UTRs (Chr4II and Chr4III) of different genes. The red bar above the figure indicates the chromosomic region with read depth values ≤1. **B**) Contigs with evidence of tandem insertions for the mCherry and SP-mCherry strains. Graphs above the contigs represent the read depth for each base of the contig.

**Additional file 5. Genotyping and genomic analysis of the PH1-**Δ**osp24 mutant and the** Δ**osp24 complemented strains. A)** To select strains where the *osp24* gene was deleted, three different PCR products were amplified. Primer combinations O11 – O12 and O13 – O14 were used to verify the insertion of the Hyg cassette into the *osp24* locus. Primer combination O9 – O10 evaluates whether the *osp24* coding sequence was deleted. **B)** To select PH1-Δosp24 complemented strains, five different PCR products were amplified. Primer combinations P5 – P6 and O9 and P10 were used to test correct insertion of the cassette into the TSI locus 1. Primer combinations P7 – P8 and P11 – P12 were used to evaluated successful recombination between the two PCR fragments and whether each transformant was homokaryotic for the transgene, respectively. Primer combination O9 – O10 evaluates presence of *osp24* in the transformed strains. Red asterisks indicate the expected PCR size bands. **C)** Top IGV screenshot shows sequencing reads aligned to the FGRAMPH1_01G15939 (*osp24*) genomic region in PH-1 and PH1-Δosp24-1. Lower IGV screenshot shows that the coding sequence of *osp24* was replaced by the hygromycin cassette. Bar graph (right) represents the average read depth values for the hygromycin cassette and the two genes (FGRAMPH1_01G15937 and FGRAMPH1_01G15941) flanking the *osp24* locus. Values above the bars are the ratio value calculated as indicated above. Error bars represent standard deviation of each average coverage value. **D)** IGV screenshots displaying genomic regions with low read coverage in the mutant strain and PH-1. Regions with low coverage were usually identified at the telomeric regions (Chr1I, Chr1IV, Chr2, Chr3II). Other regions such as 5’UTRs (Chr1II, Chr1III and Chr4I) or 3’UTRs (Chr4II) of different genes and an intergenic region (Chr3I) were identified. The red bar above the figure indicates the chromosomic region with read coverage values ≤1. Values shown in brackets in the coverage section are the count range for the bar graph.

## Data availability

The data supporting all the findings of this study are available within the paper and its supplementary data. Raw read data from the different sequenced strains are available at ENA (European Nucleotide Archive) with accession number # PRJEB64490.

## Author contributions

**MD** and **KHK** designed the project, planned the experiments, and co-wrote the manuscript. **MD** and **NK** cloned the constructs for fungal transformation, transformed the fungus and performed virulence assay in wheat ears. **MD** performed all the confocal evaluations, designed the bespoke bioinformatic pipeline and analysed the sequencing data. **AM-W** and **MG-M** developed the coleoptile infection assay. **DS** defined the sequencing platform and provided bioinformatic expertise. **MU** and **AB** identified the TSI locus 1. **MU** prepared libraries for WGS. All the authors reviewed the manuscript.

## Acknowledgements

We thank the former wheat pathogenomics bioinformatician Dr John Antoniw for the analysis that supported the identification of the micro-region. Dr George Lund (Rothamsted Research, Harpenden, UK) for advice about mapping tools. All experiments involving *F. graminearum* strain PH-1 and isogenic transformants were conducted in biological containment facilities under DEFRA licence number 101948/198285.

