## Supplementary figures and images for "Identification and functional characterisation of a locus for target site integration in *Fusarium graminearum*"

### Additional file S1

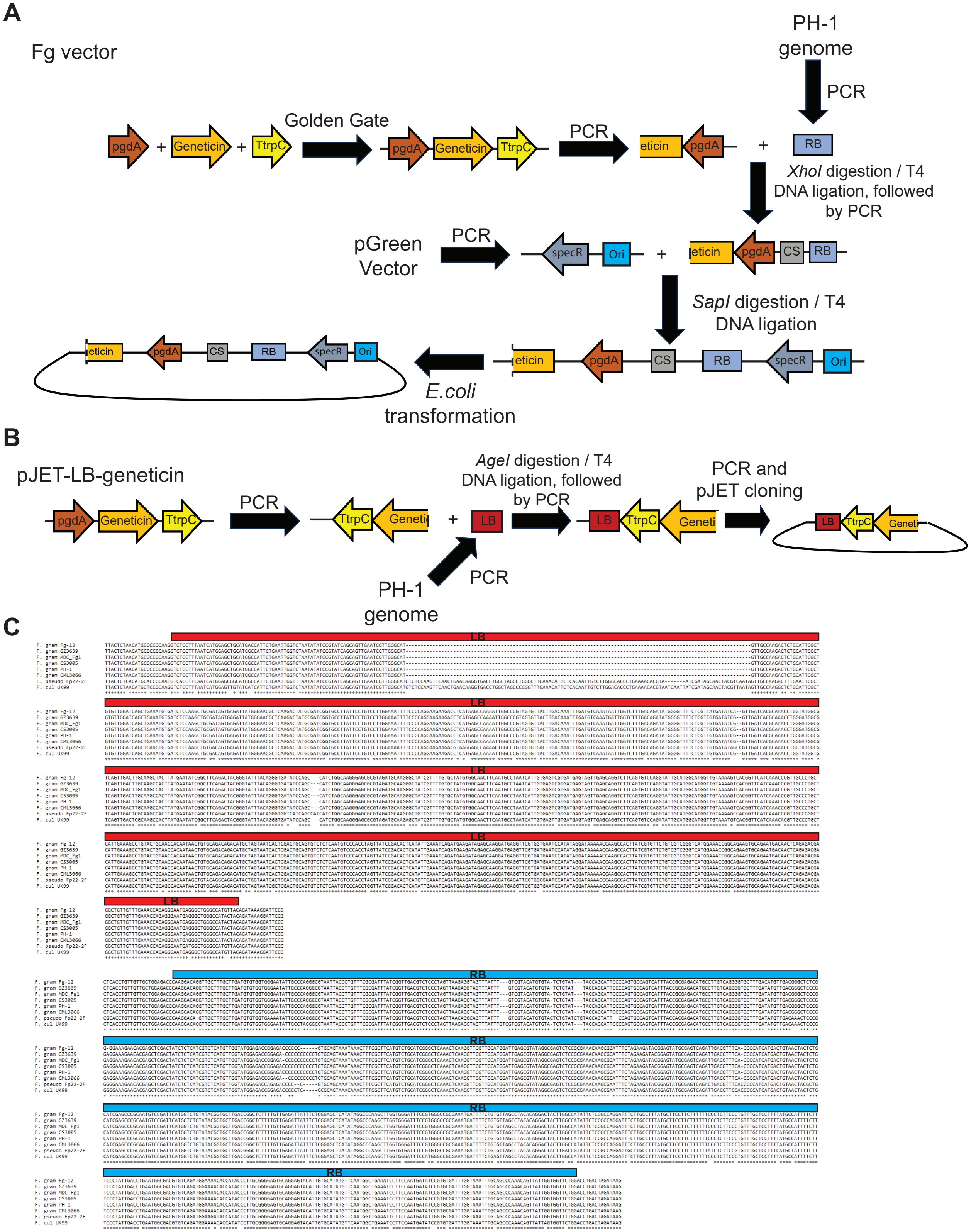

### Additional file S3

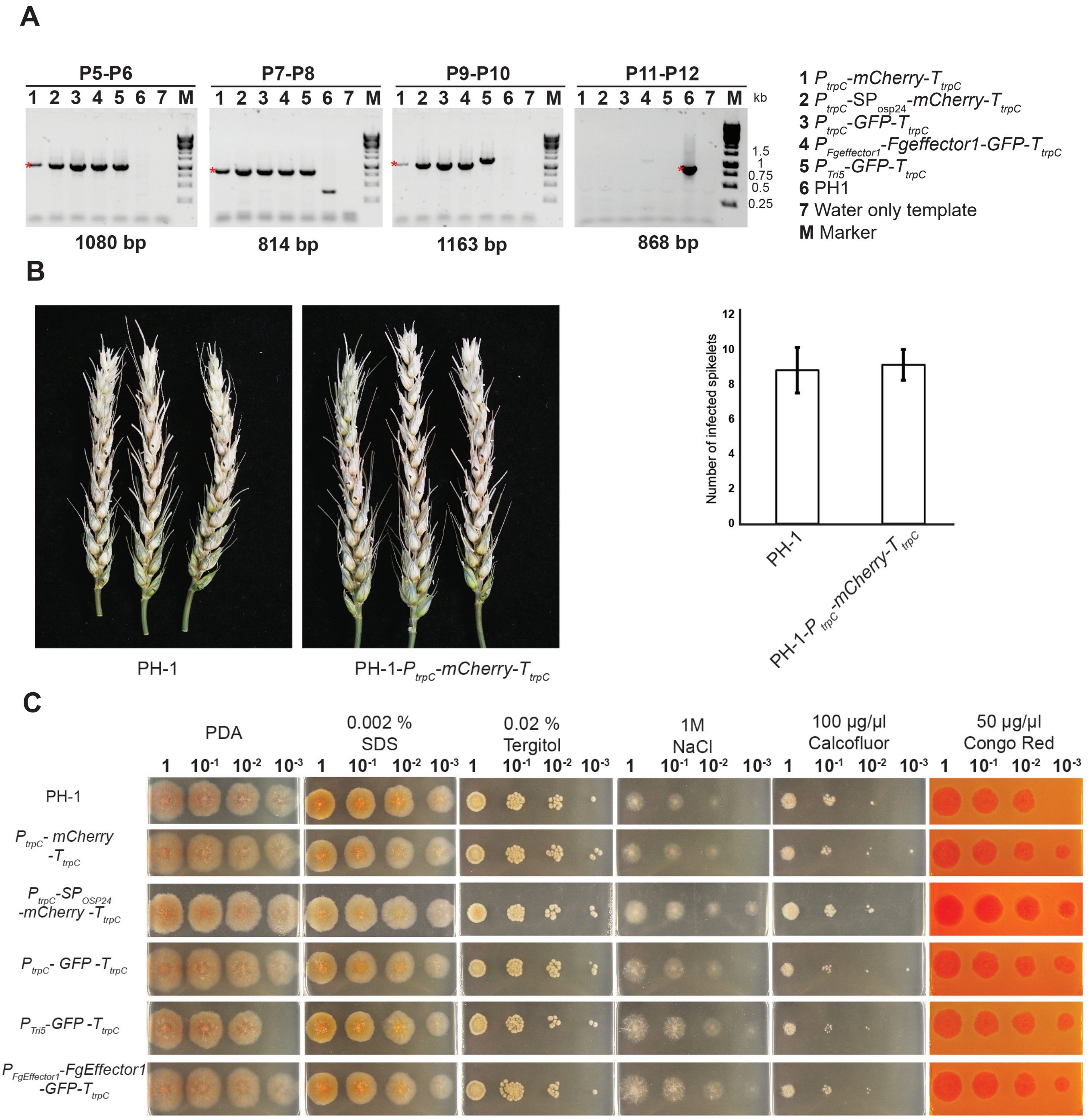

### Additional file S4

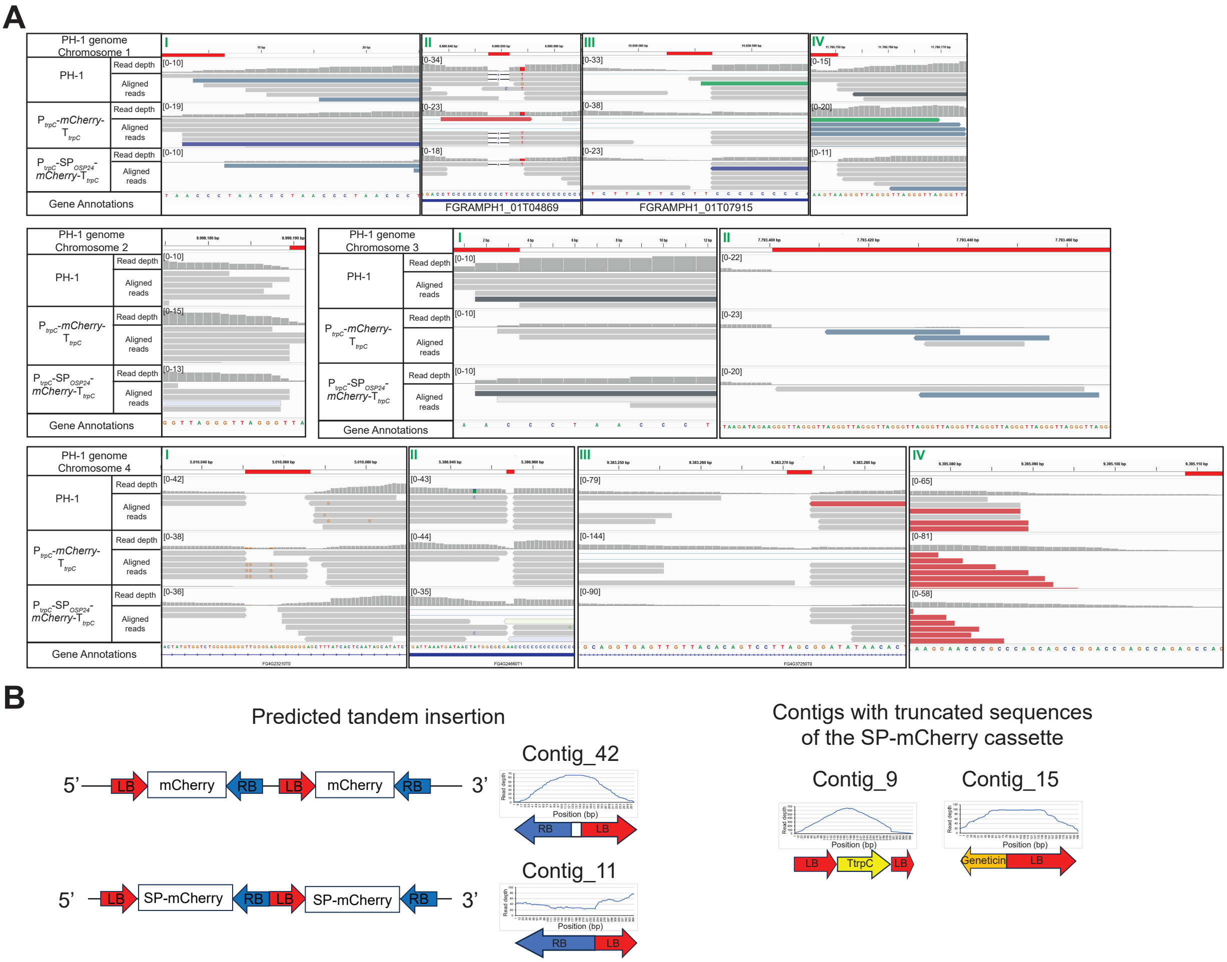

### Additional file S5

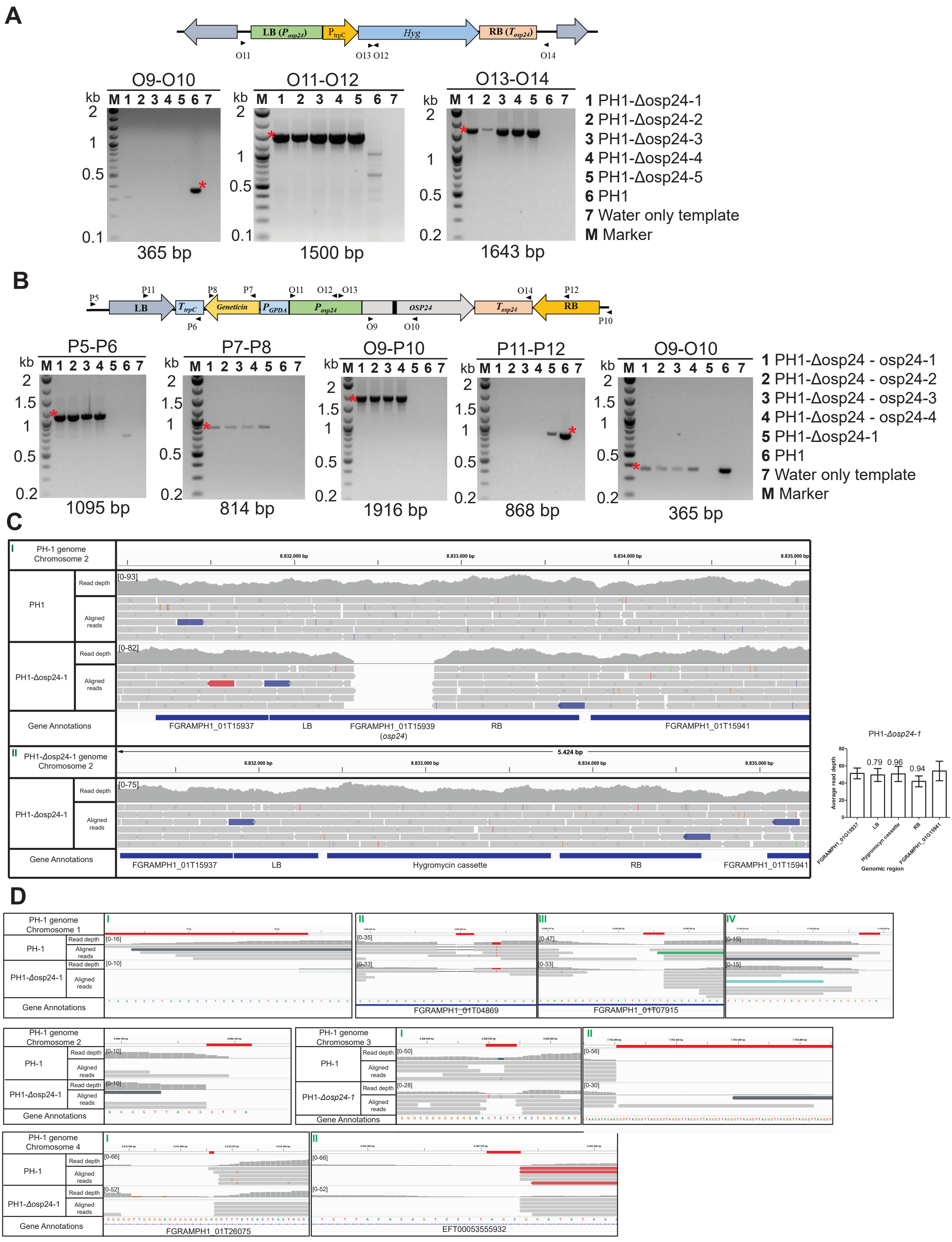
