## Additional file S2 for "Identification and functional characterisation of a locus for target site integration in *Fusarium graminearum*"

**Table 1.** List of primers used in this work

| Primer | 5'-3' sequence | Fragment amplified |
|---|---|---|
| Dest_F SapI | TATGCTCTTCATCAGGAGATCTCATGTGAG | Spec-Ori from pGreen |
| Dest_R SapI | TATGCTCTTCATGTTTAATTCCGGGGATCG | |
| FgRB_R SapI (P4)[2] | TATGCTCTTCATGACCAGAACCACCAATAACTG | RB from PH1 |
| FgRB_F XhoI | TATCTCGAGGGTCTCTACTAAGACCCAAGGACAGGTTGC | |
| pGPDApro_F | ATATGGTCTCAACCTCCTCACTCCACCATGTTGG | *Pgpda* promoter |
| pGPDApro_R | ATATGGTCTCATGTTTTTGAAGATTGGGTTCCTC | |
| TtrpC_F | ATATGGTCTCACTGCCACTTAACGTTACTGAAATC | *TtrpC* terminator |
| TtrpC_R | ATATGGTCTCATAGTGGTACCAGTTCAAGGAAGAAACAGTGC | |
| Gene_F1 (P7)[2] | ATATGGTCTCAGGCTGTATGATCGAGCAGGACGGACTCC | Geneticin gene |
| Gene_R1 (P8)[2] | ATATGGTCTCACTGATCAGAAGAATTCGTCCAACAGG | |
| Gene_R XhoI | TATCTCGAGGGTCTCAAGGTCCTCACTCCACCATGTTGG | geneticin$_{1\text{-}664}$-$P_{gpdA}$ |
| Gene_F BsaI | ATAGGTCTCACGATGTCCTGGTAGCGATCC | |
| Gene_F SapI (P3)[2] | TATGCTCTTCAACACGATGTCCTGGTAGCGATCC | geneticin$_{1\text{-}664}$-$P_{gpdA}$-RB |
| FgRB_R SapI (P4)[2] | TATGCTCTTCATGACCAGAACCACCAATAACTG | |
| TtrpC_F AgeI | ATATACCGGTGGTACCAGTTCAAGGAAGAAAC | geneticin$_{128\text{-}795}$-$T_{trpC}$ |
| Gene_R2 BsaI | TATGGTCTCACCGACCAGTCCTGTTCGTTA | |
| FgLB_F1 | ATATGGTCTCTAACAGTCTCCTTTAATCATGGAGCTGC | LB from PH1 |
| FgLB AgeI_R | ATATACCGGTTAGTATGGCCCAGCCCTC | |
| FgLB_F XhoI (P1)[2] | ATATCTCGAGGTCTCCTTTAATC | LB-TtrpC-geneticin$_{795\text{-}128}$ |
| Gene_R XhoI (P2)[2] | ATATCTCGAGCCGACCAGTCC | |
| ProTri5_F1 | ATATGGTCTCAACCTTCCTAGAACTAAGACATGG | *tri5* promoter fragment 1 |
| ProTri5_R2 | ATATGGTCTCATGTTGATGGCAAGGTTGTACTGG | |
| ProTri5_R1 | ATATGGTCTCAGTCCTTGACGTATGGACGTGCTC | *tri5* promoter fragment 2 |
| ProTri5_F2 | ATATGGTCTCAGGACTCTCTTCACGACTGTCTGG | |
| ProFgEffector1_F | ATATGGTCTCAACCTTCGGCTATCATACCATCACG | FgEffector1 promoter |
| ProFgEffector1_R | ATATGGTCTCATGTTGATGAACGTTTGAAAAAGGTAGTC | |
| PtrpC_F | ATATGGTCTCAACCTAGTCGCTGCAGGAATTCG | *trpC* promoter |
| PtprC_R | ATATGGTCTCATGTTTTGGATGCTTGGGTAGAA | |
| FgEffector1_F | ATATGGTCTCAGGCTGTATGTGCACCCAGGGACTCAAGTA | FgEffector1 gene |
| FgEffector1_R | ATATGGTCTCACTGATTTGCCAATGCCTGTTG | |
| P5 | TGCTACAGACAAAACCCGCT | LB genotyping |
| P6 and P9[1] | GCGACCTACGAGACTGAGGAATCCGCTCTTGG | |
| P10 | AGATGGACTCCCGGACATCA | Combine with P6 for RB genotyping |
| P11 | GTTCATCAAACCCGTTGCCC | To test insertion in TSI locus 1 |
| P12 | TTTGAGCCCGATGCAGACAT | |
| O1 | CATGGCGGCCGCGGGAATTCGATTAGACCATTACGGGCCTG | $P_{osp24}$ promoter |
| O2 | GTTATCGAATAGACTGTGGTGTTTGGATTC | |

| | | |
|---|---|---|
| O3 | ACCACAGTCTATTCGATAACTGATATTGAAGGAGCATTTTTT | |
| O4 | GCCGCGAATTCACTAGTGATGGATGCCTCCGCTCGAAG | *PtrpC*-HyG$_{1-761}$ |
| O5 | CATGGCGGCCGCGGGAATTCGATCGTTGCAAGACCTGCCTG | Hyg$_{296-1027}$ |
| O6 | CTTGACTCCTCTCGAGGTCGACGGTATC | |
| O7 | CGACCTCGAGAGGAGTCAAGACTGGAATC | $T_{osp24}$ terminator |
| O8 | GCCGCGAATTCACTAGTGATCACAATAGGAAAGTAAATTGAGATAG | |
| O9 | TTGATCGGTTTCCTGGGAGC | *To test osp24 presence* |
| O10 | CACAGTTTCCTCGCGTGTTG | |
| O11 | CATGATATATCACATTCTCGGCGG | *LB genotyping of* PH1-Δosp24 |
| O12 | ACTTCTCGACAGACGTCGC | |
| O13 | ACTCACCGCGACGTCTGT | *RB genotyping of* PH1-Δosp24 |
| O14 | CCAGTTAGTAGCCTGCCCAC | |
| O15 | TGGGTCTCGACCTTCTTACTCTGCAGAAATCAA | *osp24* gene fragment 1 |
| O16 | TGGGTCTCGGTCCTAGTTCTCTCTCAATTTAGTC | |
| O17 | TGGGTCTCGGGACTCGCAGGACTGGGATATA | *osp24* gene fragment 2 |
| O18 | TGGGTCTCGTAGTCACAATAGGAAAGTAAATTGAGAT | |

[1]P9 is the same primer as P6, because a primer that anneals to the $T_{trpC}$ terminator was select.

[2]Primers in brackets were also used for validation of the transformants.
